# Linking fluid-axons interactions to the macroscopic fluid transport properties of the brain

**DOI:** 10.1101/2023.01.06.523035

**Authors:** Tian Yuan, Wenbo Zhan, Daniele Dini

**Author notes:** Corresponding author (T. Yuan); (W. Zhan); (D. Dini).

## Abstract

Many brain disorders, including Alzheimer’s Disease and Parkinson’s Disease, and drug delivery procedures are linked to fluid transport in the brain; yet, while neurons are extremely soft and can be easily deformed, how the microscale channel flow interacts with the neuronal structures (especially axons) deformation and how this interactions affect the macroscale tissue function and transport properties is poorly understood. Misrepresenting these relationships may lead to the erroneous prediction of e.g. disease spread, drug delivery, and nerve injury in the brain. However, understanding fluid-neuron interactions is an outstanding challenge because the behaviours of both phases are not only dynamic but also occur at an extremely small length scale (the width of flow channel is ∼100 nm), which cannot be captured by the state-of-the-art experimental techniques. Here, by explicitly simulating dynamics of the flow and axons at the microstructural level, we, for the first time, establish the link between micromechanical tissue response to the physical laws governing the macroscopic transport property of the brain white matter. We found that interactions between axons and the interstitial flow is very strong, thus playing an essential role in the brain fluid/mass transport. Furthermore, we proposed the first anisotropic pressure-dependent permeability tensor informed by microstructural dynamics for more accurate brain modelling at the macroscale, and analysed the effect of the variation of the microstructural parameters that influence such tensor. These findings will sheds light on some unsolved issues linked to brain functions and medical treatments relying on intracerebral transport, and the mathematical model provides a framework to more realistically model the brain and design the brain-tissue-like biomaterials.

**Statement of significance:** This study reveals how neurons interact with the fluid flowing around them and how these microscale interactions affect macroscale transport behaviour of the brain tissue. The findings provide unprecedented insights into some unsolved issues linked to brain functions and medical treatments relying on intracerebral fluid transport. Furthermore, we, for the first time, established a microstructure-informed permeability tensor as a function of local hydraulic pressure and pressure gradient for the brain tissue, which inherently captures the dynamic transport property of the brain. This study is a cornerstone to advance the predicting accuracy of brain tissue transport property and neural tissue engineering.

## 1. Introduction

Brain, one of the most complex organs in nature, regulates almost all the physiological and mental activities in our bodies; yet the mechanisms behind these regulations are far from being thoroughly understood [1, 2]. The brain tissues and neuron cells are surrounded by the cerebrospinal fluid (CSF) and interstitial fluid (ISF) [3]; transport of these fluids plays a vital role in maintaining brain function and health by transferring nutrients [4], clearing waste [5], delivering drugs [6], and even directing the neuron growth [7]. Evidence has shown that some brain mechanisms (e.g. sleep [8, 9]) and brain disorders (e.g. prionlike diseases [10, 11]) are closely linked to the fluid flow in the brain.

The potential success of the emerging convection enhanced delivery (CED) method, which can physically bypass the brain-blood barrier and deliver drugs directly into the brain tissue, also depends on the understanding of transport properties of the brain [12, 13]. Other examples of the importance of mastering the laws governing fluid transport in brain for variety of applications have been systematically reported in Ref. [14]. Meanwhile, doubts have also been raised to question the role of bulk flow in transporting the interstitial solute, with some researchers suggesting that it is molecular diffusion and not convection that dominates the solute transport in the brain [15]. This has, in fact, highlighted a demand for a deeper and clearer understanding of how fluids flow in the brain.

Morphologically, the brain is a type of porous material, so fluid flow in the brain is commonly described as porous flow and governed by Darcy’s Law [16], considering the very low Reynolds number (at the level of 1E-5 even with external infusion [12]). Hydraulic permeability, which measures how easily fluids can flow through the porous domain, is the fundamental material parameter to characterise the fluid transport property of a material, but a precise evaluation of the brain permeability, especially in white matter (WM), is extremely difficult and has, at least until recently, eluded scientists. The main reasons are: (i) the micro-channel network allowing for fluid flow is anisotropic due to the directionality of the nerve fibres [17], so all components of the permeability tensor need to be determined [18]; (ii) axons are extremely soft, with Young’s modulus normally at the level of 10^2^ ∼ 10^3^ Pa [19]. Even low hydraulic pressures (e.g. 300-600 Pa) or pressure gradient (e.g. 1 Pa/*μ*m) may readily deform the axons and thus change the flow channel network, whereas the standard intracranial pressure (ICP) could be as high as 1000 ∼ 2000 Pa [20]. As anisotropy and extreme deformability are essential characteristics of the brain microstructure, unravelling their macroscale influence is not only significant to understand fluid/mass transport in the brain but also to shed light on other important research topics, e.g. the microstructural design of brain-tissue-like biomaterials [21, 22, 23, 24, 25] to mimic brain’s hydraulic properties, such as poroviscoelastic responses and fluid/mass transport property [26].

Recent advances in measuring techniques and theories have made it possible to solve these two issues at the tissue scale (as perceived at the macroscale). Examples include the directional infusion experiments into the ovine brain tissue to measure the anisotropic permeability tensor [27] and the mathematical models based on poroelastic/biphasic theory, which consider the pressure-driven deformation of the porous matrix [28, 29]. However, while our recent study has unveiled the decisive role of local microstructure on fluid transport property in the brain [18], no existing method or technique can explicitly describe the contribution of the microstructure at the axon scale and explain how the ISF acts on the axons while squeezing through the narrow network of channels bounded by the axons and available for the fluid to flow. In addition, no study has explained the mechanism behind the direction dependence of these ISF-Axon interactions and fluids transport, while some experimental evidence has been recently provided to demonstrate a clear difference in tissue responses to infusion from different directions [27]. Furthermore, due to the lack of these fundamental insights, there is no anisotropic pressure-dependent permeability tensor for brain modelling, which has largely hindered the development of accurate mathematical models for the brain tissue at the macroscale.

In this work we establish a comprehensive modelling platform for the study of fluid transport in brain tissue. This is achieved through integrating multiple models to a physically-accurate description of WM, including reconstructing the microstructure of WM from high-resolution imaging data, modelling the fluid-solid interaction (FSI) between the ISF and axons, describing the axons using a realistic visco-hyperelastic model, and explicitly considering the multibody contact between the axons driven by large deformation. We numerically reproduced the ISF-Axon interaction phenomenon in the microstructure of brain WM, which successfully unravelled the origin of the macroscale anisotropic pressure-driven transport property (permeability) of the brain WM. Furthermore, the first anisotropic pressure-dependent permeability tensor for brain modelling was established and validated using ovine brain infusion experiments. We also used geometries reconstructed using parameters obtained from high-resolution imaging of the WM microstructure to study the effect of microstructural variation on the macroscale transport property and enhance our findings. This study provides not only a more explicit and deeper understanding of how fluid/mass transport in the brain, but also a versatile tool to explore brain mechanics [30, 31].

## 2. Materials and methods

### 2.1. Geometry reconstruction

Nerve fibres (axons) constitute the WM and their directionality makes the WM resemble the anisotropic fibrous composite. As the axons are wavelike cylindrical structures, as shown in Fig. 1a [32] and Fig. 1b [33], We reconstructed the geometry as shown in Fig. 1c to represent the microstructure of WM to study the ISF-Axon interactions. We have built the dataset of WM microstructural information using FIB-SEM [34], and here, we applied the measured values to this geometry to make it as representative as possible. We chose the data from Corona Radiata (CR), where the average tortuousity (the ratio of the real length to the distance between the two ends) and diameter of the axons were 1.1 and 1.52 *μ*m, respectively. The volume fraction of the brain ECS, namely porosity, is recorded between 0.15 to 0.3, while an average value of 0.2 has been widely acknowledged [15, 35]. Therefore, 0.2 was chosen as the porosity in this model.

**Figure 1:**
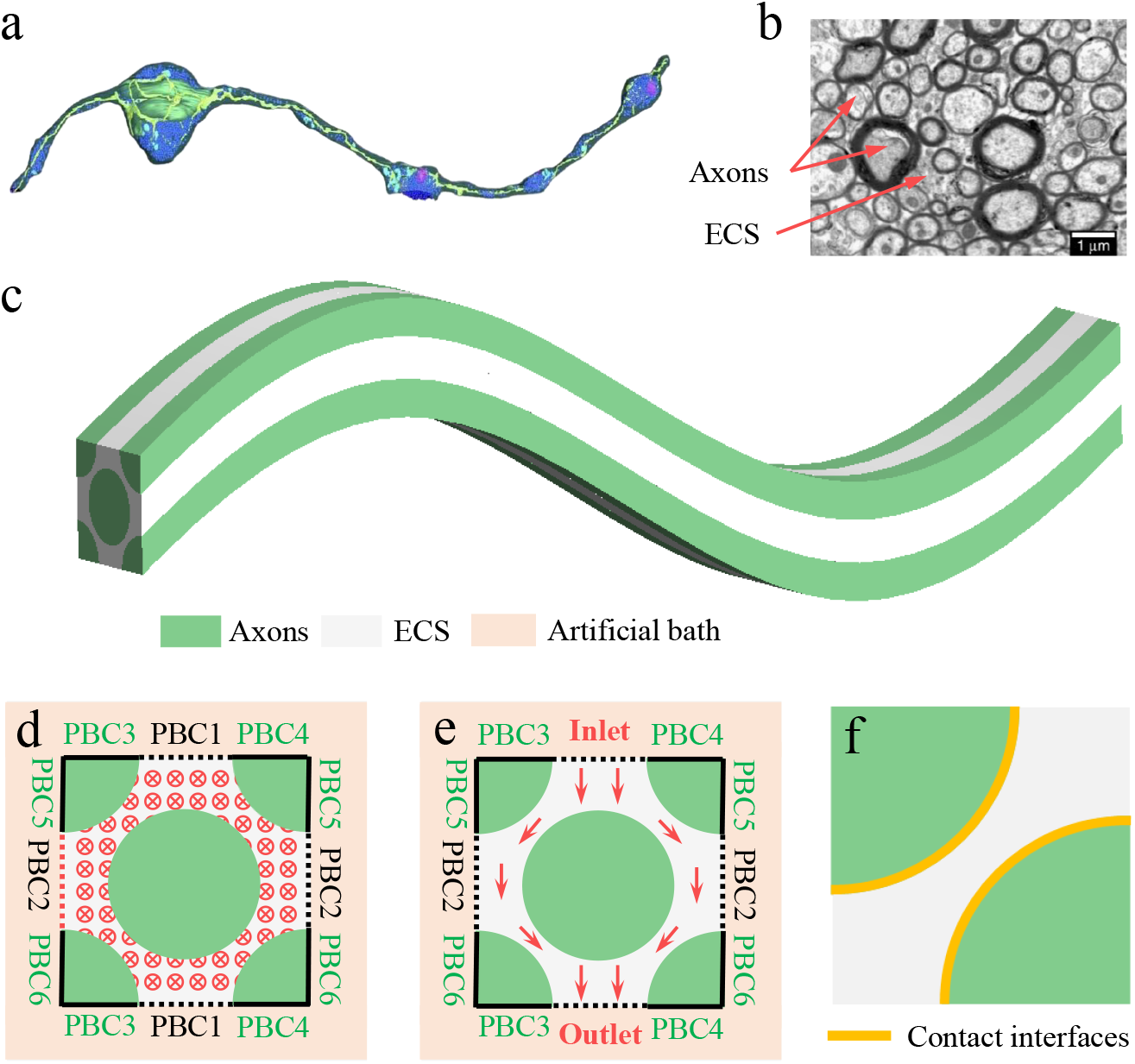
3D geometry and boundary conditions. **a**, Typical structure of axons from Focused Ion-Beam Scanning Electron Microscopy (FIB-SEM) image. The dark blue bulges consist of synaptic vesicles; their stiffness is negligible compared with the axons, so they are excluded in the geometry [32]. **b**, Typical cross-sectional configuration of white matter (WM) [33]. **c**, Reconstructed microstructure of WM. **d**, Boundary conditions of parallel flow in the cross-sectional view. **e**, Boundary conditions of perpendicular flow in the cross-sectional view. The artificial bath allows the outward deformation of axons. **f**, Illustration of solid contact between axons.

This system enables us to explicitly model the ISF flow both parallel and perpendicular to the axons with the setups showing in Fig. 1d and 1e, respectively. Periodic boundary conditions were imposed on boundaries of both axons and extracellular space (ECS) to reduce and eliminate the possible size and boundary effects. An artificial bath outside the core model was built to allow for the outward displacement of the axons driven by the hydraulic pressure. In both flow scenarios, axons may be forced to contact with each other, so the model must include algorithms that deal with contact between axons, as shown in Fig. 1f. It is worth mentioning that the ECS is not composed exclusively of ISF; there are also proteoglycans, hyaluronan, and other substances in the ECS and form the ECM. However, as these substances can absorb and release water molecules quickly [14] and ECM is far softer than axons [36], we did not explicitly model all the ECM components here. In order to simplify the description of the domain to be simulated and significantly reduce the model complexity, and in line with what has been suggested by other researchers [37, 38], we treated the WM as transversely isotropic [39]; this implies that the axons and ISF behaviours on the plane perpendicular to the main fibre trajectory is direction-independent.

The real microstructure of WM is more complex than this reconstructed geometry, but too much geometrical complexity would add irreparable divergence to the numerical analysis, especially when considering the high nonlinear terms (i.e., FSI and multibody contact). Therefore, after carefully balancing the geometrical authenticity and numerical operability, this geometric model was finally chosen.

### 2.2. Mathematical models for ISF-Axon interactions

#### Fluid Phase

The ISF was modelled as an isothermal incompressible Newtonian fluid that is governed by the Navier–Stokes (NS) equations. The inertial terms can be neglected because of the low Reynolds number. If also neglecting gravity, the ISF flow can be described by Eqs. (1) and (2).

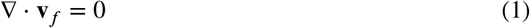

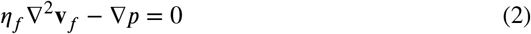

where **v**_*f*_ is the flow velocity, *η*_*f*_ is the dynamic viscosity of ISF, *p* is the local hydraulic pressure.

#### Solid Phase

Various constitutive models have been developed to describe the axonal mechanical behaviours, including linear elastic model [40], viscoelastic model (time- and rate-dependent response) [41], and hyperelastic model (sharp increase in stress with strain under large deformation due to the contractility generated by the molecular motors) [42]. To cover all these types of axonal mechanical behaviours, the axons were, for the first time, modelled as a visco-hyperelastic material in this study. As the visco-hyperelastic behaviour results from a combination of hyperelatic and viscoelastic responses, the total stress on the axon is the sum of these individual components [43]:

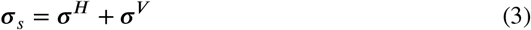

where ***σ***_*s*_ is the total stress of the solid phase (axons), ***σ***^*H*^ is the hyperelastic stress, and ***σ***^*V*^ is the viscoelastic stress. The hyperelastic response was described by the Ogden model [44], and the strain energy density function is:

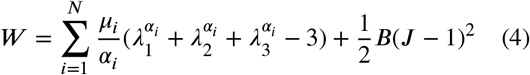

where *N* is the order of the model, which was 1 for axons; *μ*_*i*_ and *α*_*i*_ are material constants; *λ*_*j*_(*j* = 1, 2, 3) are the principal stretches; *B* is bulk modulus to capture the compressibility of axons; 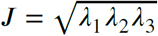 is the elastic volumetric deformation.

The Cauchy stress can be then written as:

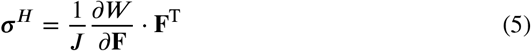

As the axonal displacement/deformation is driven by the hydraulic pressure, and creep dominates the viscoelastic response, the Kelvin–Voigt model was chosen to describe the axonal viscoelastic behaviour. The viscoelastic component of the stress can be therefore written as Eq. 6.

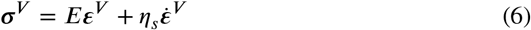

where *E* is the elastic modulus; *η*_*s*_ is the viscosity of the axons; ***ε***^*V*^ is the viscoelastic component of strain.

When a suddenly stress is applied, e.g., *σ*_0_, the strain history would follow:

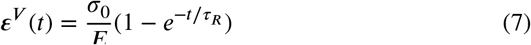

where *t* is time and *τ*_*R*_ = *η*_*s*_/*E* is the retardation time.

#### Fluid-solid interaction (FSI)

Two-way coupling scheme was adopted to describe the FSI between the ISF and axons (Eqs. 8, 9, 10), which means that the axon deformation and ISF flow status were updated in every time-step. This enables us to consider large deformation of the axons and achieve higher precision.

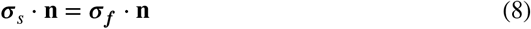

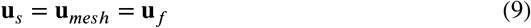

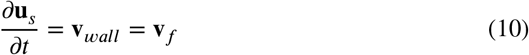

where **n** is the normal direction, **u** is the displacement, **v** is the velocity. The subscripts *s, mesh, f* denote solid phase, mesh, and fluid phase, respectively.

#### Contact between the axons

To take account gap closure due to contact between axons, contact interfaces were assigned to each pair of axon walls where contact may occur. Penalty formulation [45] was adopted to capture the multibody contact and no friction/adhesion was considered.

### 2.3. Brain infusion experiments

Fig. 2 briefly summarises the infusion experiment conducted using fresh ovine brain samples to measure the permeability. Samples were obtained from the CR region, where the axons’ directionality can be easily identified. The brains were transversely (Fig. 2b) and longitudinally (Fig. 2c) sliced to obtain the cylindrical parallel (Fig. 2d) and perpendicular (Fig. 2e) samples. Axial infusions with different pressure were performed via a syringe needle (BD MicrolanceTM; stainless steel; 30G × 1/2”; 0.3 ×13 mm). The infusion pressure (P) and flow rate (Q) were recorded for the equivalent permeability calculation, as illustrated by Fig. 2f. See Ref. [27] for more details.

**Figure 2:**
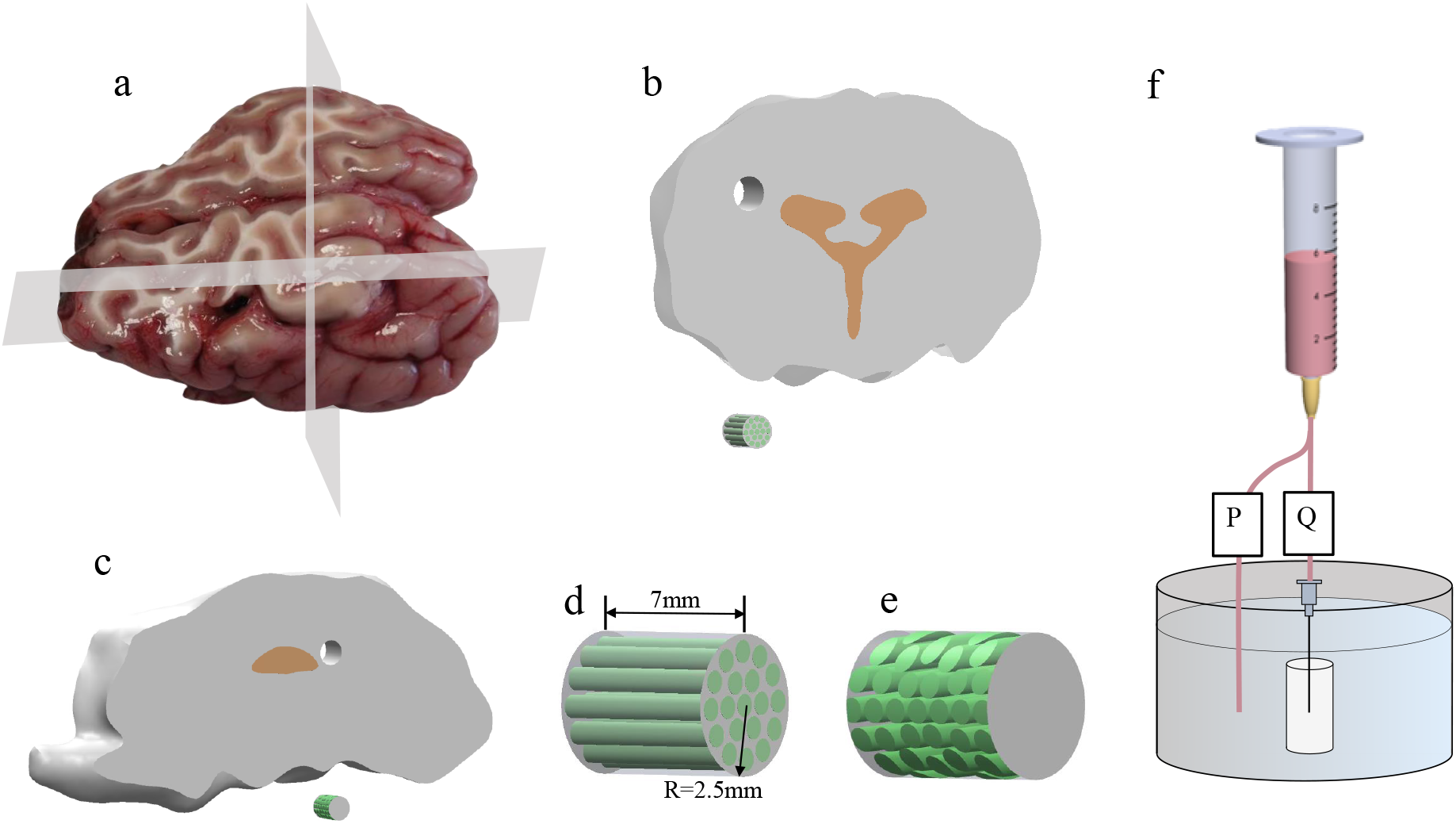
Infusion experiment with ovine brain WM samples. **a**, Fresh ovine brain was transversely and longitudinally sliced to expose the WM. **b**, Schematic representation of transverse sampling. **c**, Schematic representation of longitudinal sampling. The yellow part is the ventricle. **d**, The parallel WM sample from Corona Radiata (CR). The green structures show the orientations of the axons in the sample. **e**, The perpendicular WM sample from CR. **f**, The infusion protocol. The cylindrical samples were immersed in a PBS bath, and all infusions were along the axis of the cylindrical samples. Q denotes the flow rate sensor and the pressure sensor (P) measures the pressure difference between the syringe and the bath.

### 2.4. Tissue sample simulations

Simulations with the ovine brain sample were conducted twice, both of which adopted Darcy’s Law but with different formats and permeability tensors.

Step 1: as will be mentioned in section 3.1, the local pressure and pressure gradient change simultaneously in the real situation, we thus assigned the median values of the measured permeability tensor (*k*_∥_ = 1.5 × 10^−16^ m^2^, *k*_⊥_ = 0.7 × 10^−16^ m^2^) [27] to obtain the matching pairs of pressure and pressure gradient (Eq. 11), which were used to study the realistic coupling effect of pressure and pressure gradient (Fig. 5). Note that on this step, the pressure-dependent permeability tensor had not been obtained so that the tissue samples were treated as homogeneous. This approximation helps to obtain the coupled pairs of pressure and pressure drop, and would not change the quantitative results in terms of the effect that pressure and pressure gradient has on permeability.

**Figure 3:**
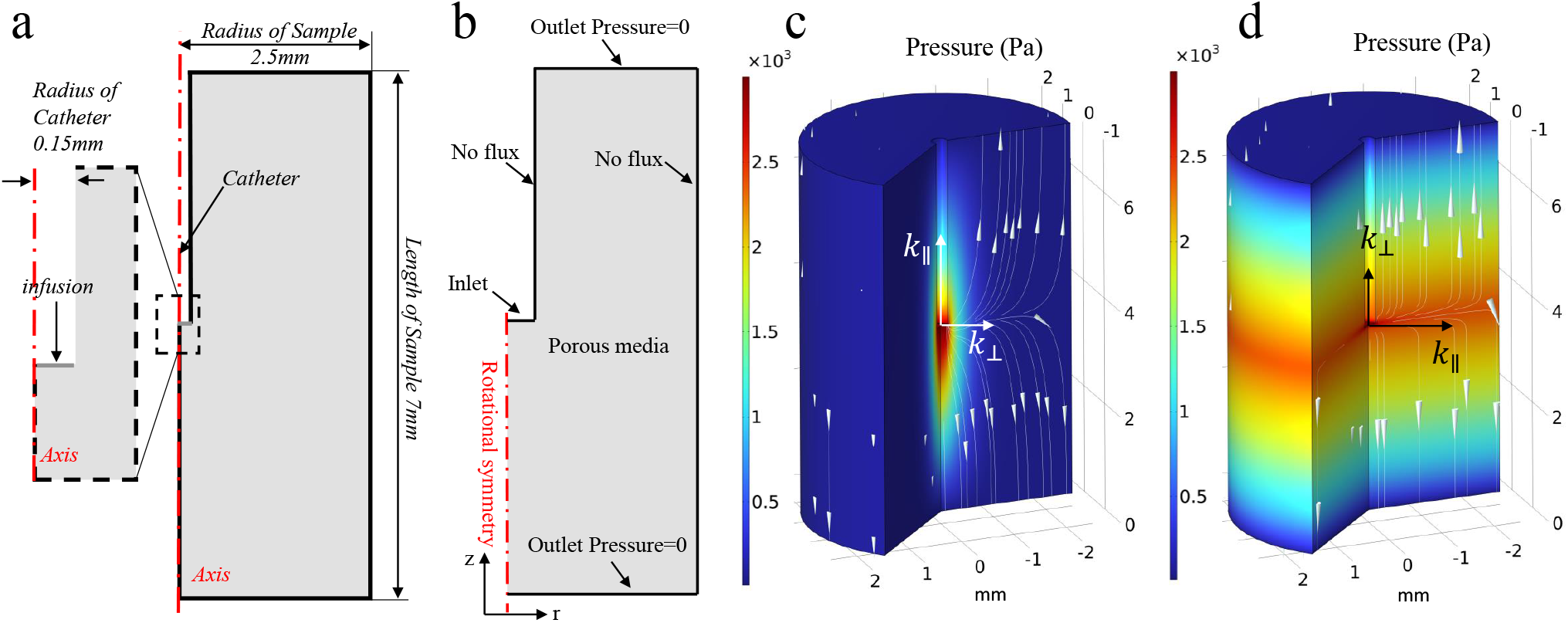
Geometry, boundary conditions, and simulation results of tissue infusion experiments. **a**, Geometric model of the ovine brain tissue sample. Since the tissue samples were cylindrical (Fig. 2d and 2e), a 2D axisymmetric model was built to represent the tissue samples. The cylindrical hole in the middle is the needle penetrating along the axis. **b**, Boundary conditions of the tissue sample. At the tip of the needle is the flow inlet boundary, and the top and bottom surfaces are the flow outlet boundaries. A cylindrical and transparent plastic tube was outside the tissue sample to support the sample so that the flow cannot pass through this surface. No flux boundary condition was thus assigned to the outer surface of the sample. The same condition exists on the surface contacting with the needle wall, so No flux boundary condition was also assigned to this surface. **c**, Simulation results of the parallel infusion. **d**, Simulation results of the perpendicular infusion. The colour legend denotes the hydraulic pressure distribution and the streamlines represent the flow direction.

**Figure 4:**
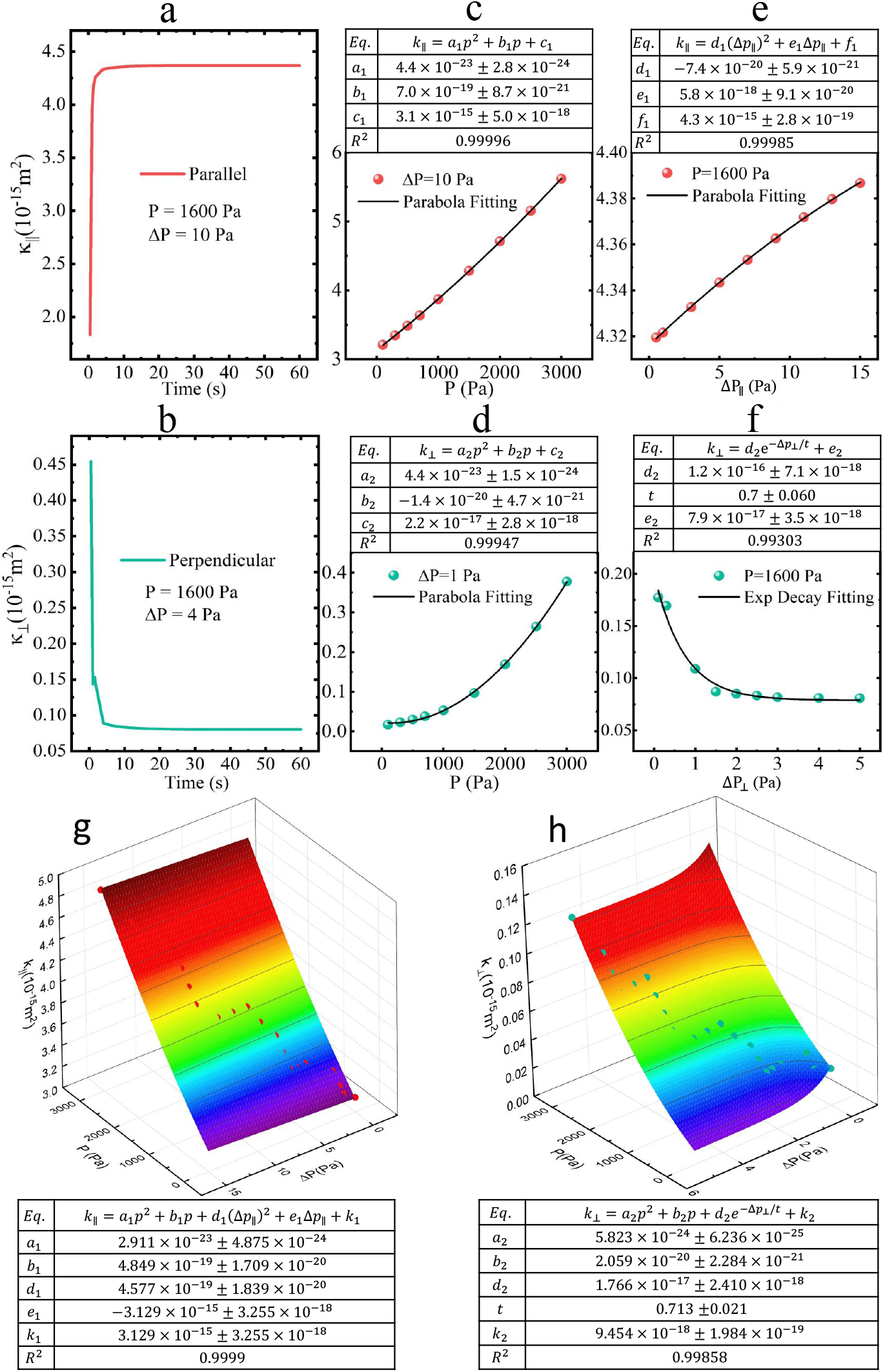
Derivation of permeability tensor. **a**, Time dependence of permeability in the parallel direction. **b**, Time dependence of permeability in the perpendicular direction. **c**, Pressure dependence of permeability in the parallel direction. **d**, Pressure dependence of permeability in the perpendicular direction. **e**, Pressure drop dependence of permeability in the parallel direction. **f**, Pressure drop dependence of permeability in the perpendicular direction. **g**, Pressure and pressure drop dependent permeability in parallel direction. **h**, Pressure and pressure drop dependent permeability in perpendicular direction. The boxes over the fitted curves show the fitting equations, values of coefficients and the *R*^2^ of each fitting result. The red and green colours denote the data in parallel and perpendicular directions, respectively.

**Figure 5:**
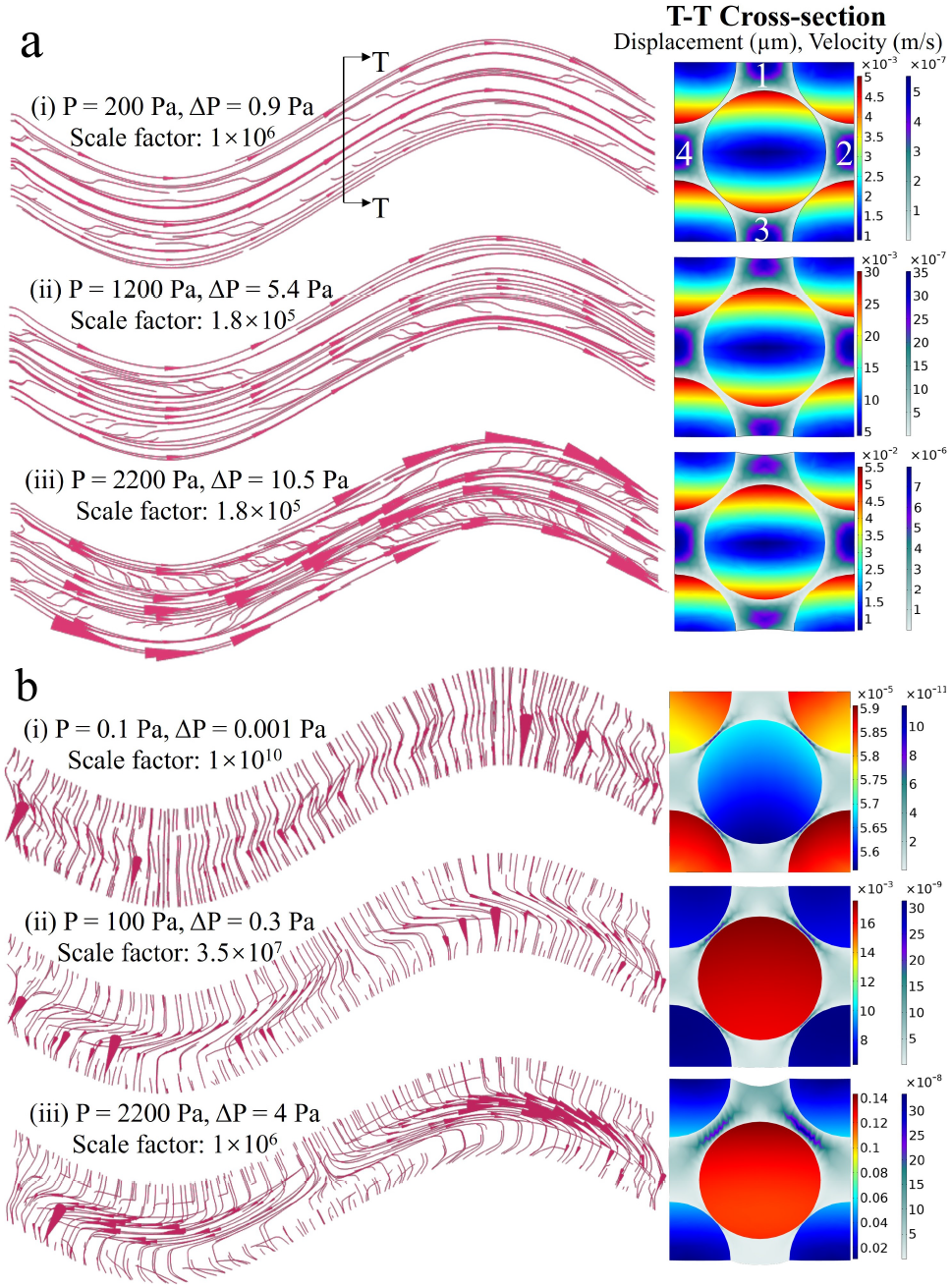
Fluid-solid interaction between the ISF and axons. **a**, The parallel flow status and axonal deformation. **b**, The perpendicular flow status and axonal deformation. The streamlines on the left-hand side figs denote the flow direction. The cone size of the streamlines is proportional to the flow velocity magnitude and the scale factors are given above the streamlines. The figs on the right-hand side show the flow velocity and axonal deformation in the middle transverse section.

Step 2: after obtaining the pressure-dependent permeability tensor with the microstructural simulations and assigning it to the tissue sample, Eq. 11 was used again to calculate the realistic flow field in the tissue sample. Eq. (12) was then used to calculate the equivalent global permeability tensor (*K*) of the tissue sample and compare it with the measured values (Fig. 6).

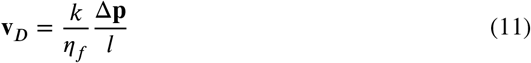

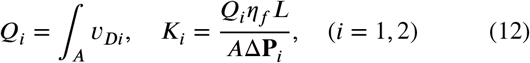

where **v**_*D*_ is the local Darcy velocity tensor; *k* is the local permeability tensor (note that it was assumed homogeneous on Step 1 and was pressure-dependent on Step 2); Δ**p** is the local pressure drop tensor; *l* is the length of the finite element; *Q*_*i*_ (SI unit: m^3^/s) is the fluid discharge rate in the sample; *A* is the cross-sectional area of the sample; *v*_*Di*_ is the Darcy velocity on the outlet surfaces of the sample; *K*_*i*_ is the sample equivalent permeability; *L* is the length of the tissue sample; Δ**P**_*i*_ is the pressure drop along the sample; components 1 and 2 denote the parallel and perpendicular directions, respectively.

**Figure 6:**
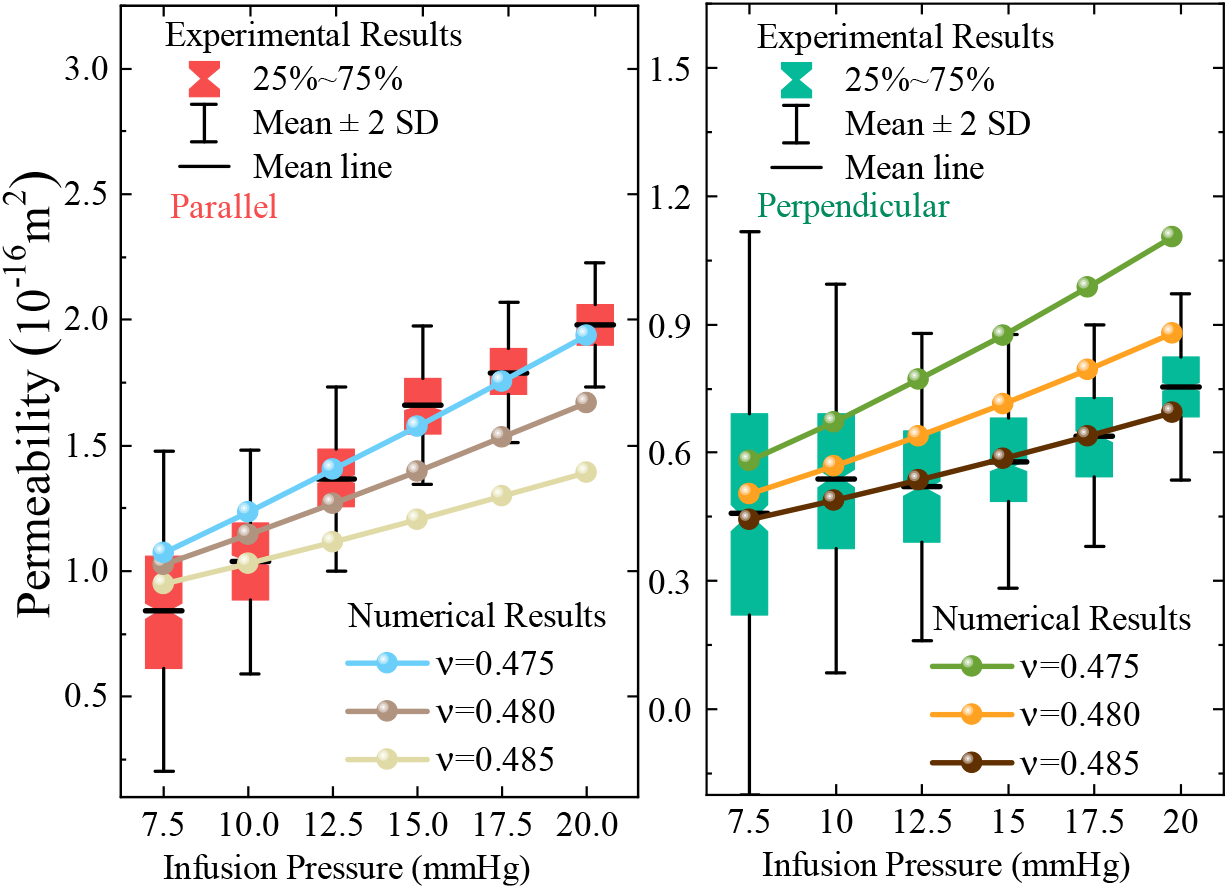
Comparison of infusion pressure-equivalent permeability relationship between experimental and simulation results. The left-hand side figure shows the parallel results and the right-hand side figure shows the perpendicular results.

The model geometry was directly built from the tissue samples (Figs. 2d and 2e), as shown in Fig. 3a. Fig. 3b shows the boundary conditions for the simulations. Fig. 3c and 3d present the representative simulation results.

### 2.5. Finite element implementation

All the simulations were conducted with COMSOL Multiphysics [45] platform and the values of relevant parameters are summarised in Table 1. Both of the microscopic model (axon model) and macroscopic model (brain tissue sample) have passed mesh sensitivity tests.

**Table 1.**
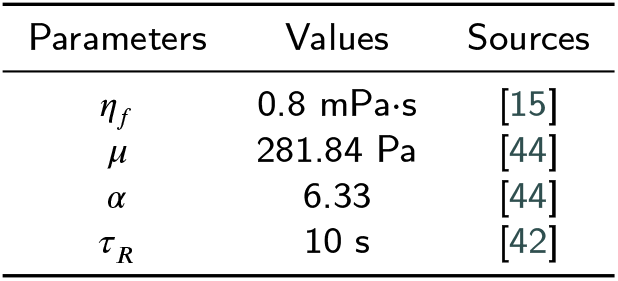
Parameters and values for the simulations Parameters Values Sources

## 3. Results

### 3.1. Permeability tensor

Given their visco-hyperelastic constitutive material behaviour [42, 44], the axons underwent dynamic displacement/deformation under a transient load, but would reach equilibrium after 10 ∼ 20 s, as shown in Fig. 4a and 4b. Movies S1 and S2 also show these transient deformation processes.

Both local pressure and pressure gradients are responsible for the deformation of the axons, thus changing the local permeability; yet these two parameters are fully coupled and change simultaneously in the real situation. As a result, their individual effects on the local permeability tensor are currently unknown. To fill this gap, we used different values of inlet pressure and pressure drop to conduct a parametric sweep. The range of pressure (0 ∼ 3000 Pa) were chosen based on the tissue infusion experiments that will be described later [27], and the range of corresponding pressure gradient (0 ∼ 15 Pa in parallel and 0 ∼ 5 Pa in perpendicular) were estimated by the method in Step 1 of Section 2.4.

Results show that parallel permeability grows almost linearly with the local pressure (Fig. 4c) while the perpendicular permeability increases parabolically with the local pressure (Fig. 4d), which implies that the perpendicular permeability is more sensitive to higher local pressure. Although a linear function can describe the pressure sensitivity - parallel permeability relationship, we found that there still exists a slight non-linearity, as demonstrated in the following section. Similar trends were observed in the perpendicular direction, so a parabolic function was used to describe the relationship between local pressure and permeability in both directions. In contrast, permeability varies very differently to pressure gradient in the two flow directions. The parallel permeability shows a parabolic increase against the pressure drop (Fig. 4e), while the perpendicular permeability shows exponential decay with the pressure drop (Fig. 4f). Thus, we can write pressure - pressure drop - permeability relationships (*k* = *f* (*p*, Δ*p*)) as follows:

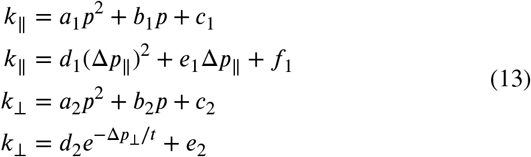

where *k*_∥_ and *k*_⊥_ are the parallel and perpendicular components of the local permeability tensor, respectively; *p* is the local pressure; Δ_*p*∥_ and Δ_*p*⊥_ are the local pressure drops in parallel and perpendicular directions, respectively; *a*_1_ ∼ *f*_1_ are the coefficients of the parallel component, and *a*_2_ ∼ *e*_2_, *t* are the coefficients of the perpendicular component.

By combining the *p* terms and Δ*p* terms in a first approximation, we can rewrite the permeability tensor as follows:

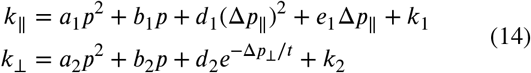

Next, we used these two equations to fit the *p* − Δ*p* − *k* data points obtained from the parametric sweep. As shown in Fig. 4g and 4h, both components of the permeability tensor fit the data well, with the *R*^2^ above 0.99. Note that this figure represents only one set of geometric parameters. More results are available in Section 3.4.

### 3.2. Flow status and axon deformation

We now turn to the dynamic flow and axonal behaviours to shed light on the origins of the links between the permeability tensor components, applied pressure and pressure gradients. Fig. 5 shows the flow status (left-hand side Figs) and axons’ deformation (right-hand side Figs) under different combinations of pressure and pressure drop. See Section 2.4-Step 1 for how these pairs were chosen. As mentioned above, the local pressure and pressure gradient in real situation are changing simultaneously and cannot be decoupled, so we mainly present the coupled effect of these two parameters here. Movies S3 ∼ S6 also show the individual effects of local pressure and pressure gradients variations within a physiological range of interest captured by parametric sweeps using the same values as the parametric sweep in the Section 3.1.

Fig. 5a shows the results of parallel flow. The streamlines denote the flow direction, the cone size is proportional to the flow velocity magnitude (see the scale factors above the streamlines), and the density of the streamlines indicates the local flow rate. The right-hand side figs also provide the value of flow velocity on the middle transverse section. Under low pressure, the streamlines are relatively sparse and gathering in the positions where gaps between axons are large (areas 1 ∼ 4 on the contour). Less transverse flows develop. The reason is that the diagonal gaps between axons is initially so narrow that flowing through them is difficult, thus the majority of fluid can only flow along the axons and gather in areas 1 ∼ 4. With the increase of the local pressure, the streamlines become denser, implying higher flow rates. More transverse flow across the axons also develops. These are due to the pressure-induced axon shrinkage, which not only provides more spaces for the flow but also opens diagonal gaps. In addition, the higher parallel flow rate and momentum straightens the axons (see Movie S3) and thus further removes the parallel resistance originated from the channel diversion. All these axonal deformations contribute to the aforementioned nonlinear increase of permeability with local pressure (Fig. 4c). Regarding the effect of pressure gradient, a higher pressure drop leads to a faster parallel flow and straighter flow pathways, thus increasing the permeability. This response, however, would gradually reach a critical point when axons reach their maximum elongation. Movie S4 also shows that this factor, in fact, has a very limited effect on the axon behaviour and channel deformation within the studied range. Consequently, Fig. 4e shows that the local permeability increases with pressure drop, but the change is limited and within 2% of the zero pressure gradient value. In summary, the above phenomenon-based analyses indicate that the parallel transport property of WM is dominated by local pressure.

Fig. 5b shows the results of perpendicular flow. Compared with the parallel flow case, the change of the applied pressure leads to more drastic changes of the flow status. Under low pressure, the flow is transverse to the axonal structures and steady. However, even with relatively small increments of the applied pressure, e.g. 100 Pa, the majority of the flow is diverted along the axons. This is accentuated when higher pressure is applied as also graphically shown by the density of the flow vectors in Fig. 5b*(iii)*. The main reason is that a higher local pressure normally develops with a higher pressure drop across the domain; this will bend the axons and close the lower diagonal gaps between the axons, as shown in Fig. 5b *right*. Consequently, the transverse flow pathway will be gradually closed, thus turning the flow to the parallel direction. This decreases the perpendicular permeability. However, the higher local pressure also compresses the axons and gives more space to the perpendicular flow at regimes where no axon contact occurs (see Movie S5). This means that the higher pressure may, in some cases, also lead to higher perpendicular permeability. Therefore, pressure and pressure gradient jointly determine the perpendicular permeability of WM. Movie S6 shows that a pressure gradient threshold exists, beyond which the axon gaps have been completely closed, so no more obvious permeability decrease can occur. This explains the the exponential decay of the perpendicular permeability with pressure drop shown in Fig. 4f.

### 3.3. Comparison with experimental results

The newly presented mathematical formulation of the pressure-dependent permeability tensor will now be tested by assigning this permeability tensor to ovine brain WM. This will enable us to verify that the proposed method can predict the pressure dependence reported in the experimental observations obtained using localised infusion experiments [27].

The pressure-permeability relationships are presented by the box charts in Fig. 6. It shows that the permeability tensor in both directions increases with the infusion pressure. In addition, the parallel component increases faster than the perpendicular component, indicating that the parallel permeability is more sensitive to the local pressure growth. These can be now explained by the trade-off between microstructural responses and the local pressure and pressure gradient. As analysed above, the parallel permeability increases with both the local pressure and pressure gradient (also as shown quantitatively in Fig. 4c and 4e). In contrast, the perpendicular permeability increases with the local pressure (Fig. 4d) but decreases with the pressure gradient (Fig. 4f).

The comparison between the simulation and experimental results of the ovine brain WM samples shown in Fig. demonstrates the importance of capturing the pressure-dependent nature of permeability in the proposed formulation of the permeability tensor. In state-of-the-art models, axons are widely regarded and treated as nearly incompressible [46, 47] - a material property that is defined by the value of Poisson’s ratio ranging from 0.475 to 0.499 in solid mechanics modelling [48, 49]. As no exact Poisson’s ratio of axons has been measured and reported in the open literature, many works thus set the Poisson’s ratio to be above 0.49 [50], or alternatively assign a high bulk modulus [51]. However, our results show that a value around 0.48 better captures the tissue response to deformation (Fig. 6). Our results also suggest that the pressure-dependent permeability of WM is highly sensitive to axons’ Poisson’s ratio and thus, to some extent, highlighting the importance of precise measurements to determine axons’ materials properties.

### 3.4. Microstructural dependence of the permeability tensor

The model in Fig. 1 is equipped with representativeness, but may still lack generality. This limitation was mitigated by investigating the microstructural dependence of the results through a comprehensive parameter study using other representative volumes characterised by different geometric values, i.e., axonal diameter, porosity, and tortuosity. See Table 2 for the parameters and values. Note that all these values are within our measured range [34] and other published data [52]. This study enabled us to capture the variation of our findings as a function of local microstructural features.

**Table 2.**
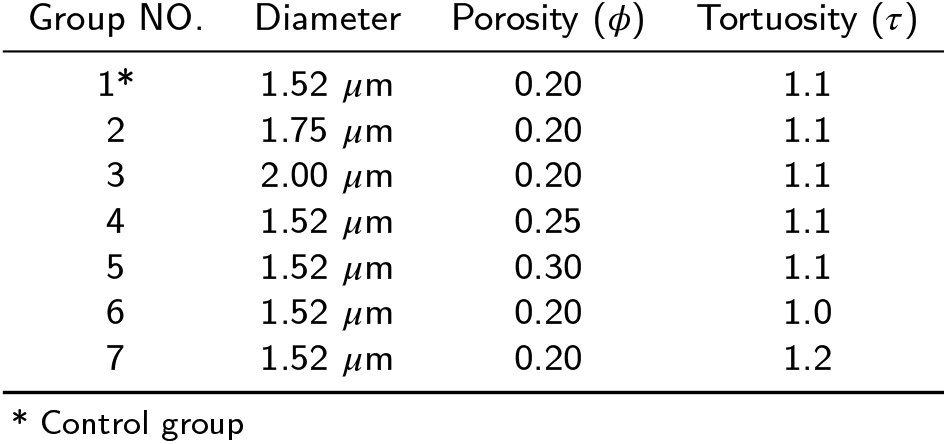
Parameters and values of the parametric study

| Group NO. | Diameter | Porosity ($\phi$) | Tortuosity ($\tau$) |
| --- | --- | --- | --- |
| 1* | 1.52 $\mu\text{m}$ | 0.20 | 1.1 |
| 2 | 1.75 $\mu\text{m}$ | 0.20 | 1.1 |
| 3 | 2.00 $\mu\text{m}$ | 0.20 | 1.1 |
| 4 | 1.52 $\mu\text{m}$ | 0.25 | 1.1 |
| 5 | 1.52 $\mu\text{m}$ | 0.30 | 1.1 |
| 6 | 1.52 $\mu\text{m}$ | 0.20 | 1.0 |
| 7 | 1.52 $\mu\text{m}$ | 0.20 | 1.2 |

\* Control group

The results are presented in Figs. 7. Overall, the varying values do not change the relationship between infusion pressure and the equivalent permeability tensor, which indicate the global validity of the established relationships for brain WM. However, the results also clearly show that small fluctuation of these values may lead to significant changes in the permeability tensor magnitude. Porosity has the most important effect on not only the magnitude of permeability tensor, but also the pressure-permeability relationship. Higher porosity leads to higher permeability magnitude and a sharper pressure-permeability curve (Fig. 7b). In the current representation, porosity is directly linked to the axon diameter, and the effects of axon diameter are similar to that of porosity (Fig. 7a). It is worth mentioning that while porosity keeps the same, larger axon diameter also leads to larger ECS and gaps between axons. According to scaling rule [53], permeability is proportional to the square of gap size, so larger axons lead to higher permeability. In terms of tortuosity, larger tortuosity brings larger resistance to the flow, thus declining permeability. Compared with axon diameter and porosity, the effect of tortuosity is less important, especially under low infusion pressure (Fig. 7c).

**Figure 7:**
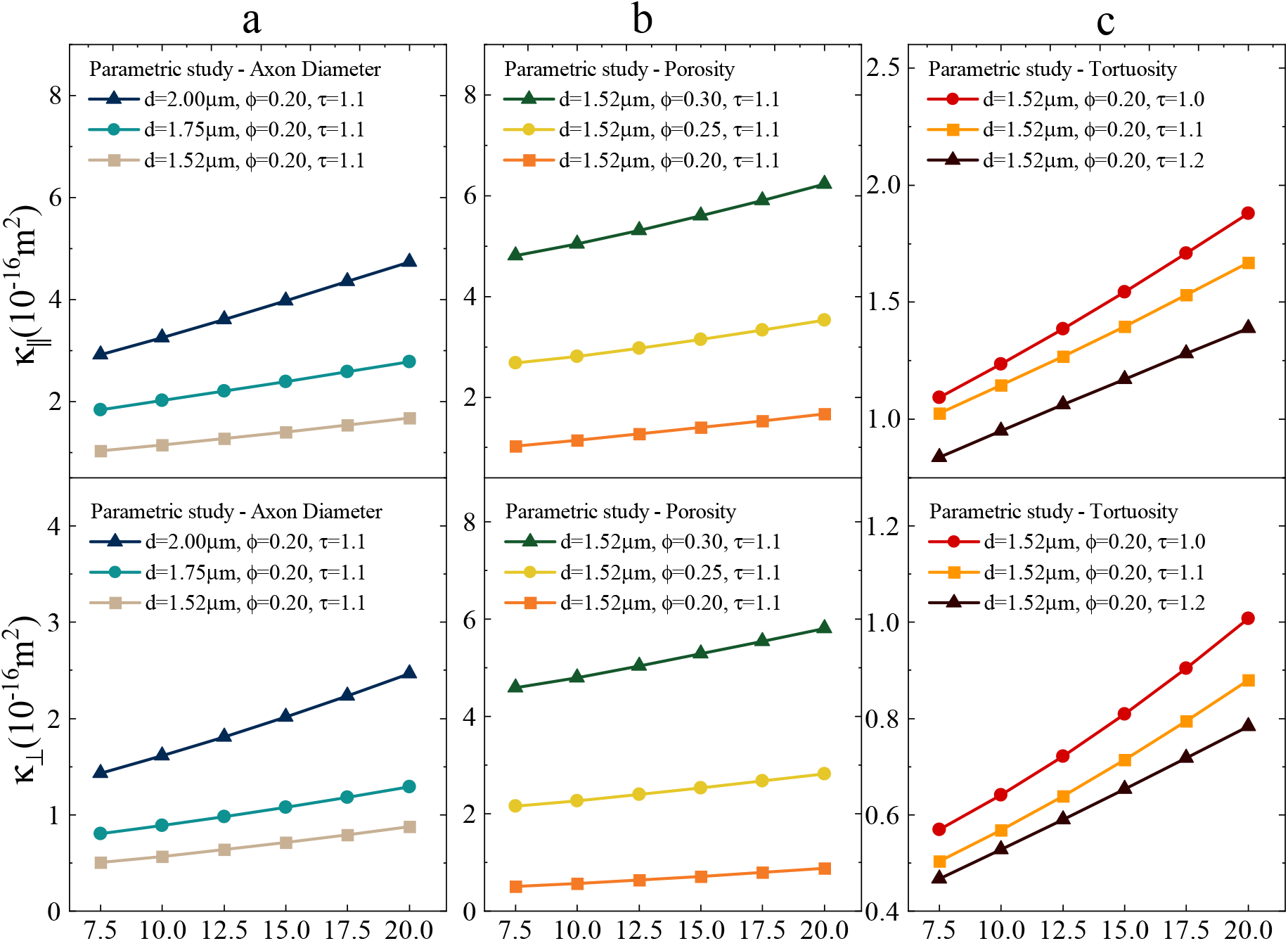
Microstructural dependence of the permeability tensor. **a**, Effect of axon diameter on the pressure-permeability relationship of WM. **b**, Effect of porosity on the pressure-permeability relationship of WM. **c**, Effect of axon tortuosity on the pressure-permeability relationship of WM.

## 4. Discussion

While the impact of fluid transport on brain functions health and has gained growing attention and substantial progress has been made to unravel the complexity that governs such phenomenon [14], a fundamental understanding of how fluids transport in the brain micro-network is still lacking. In particular, the effect that hydraulic pressure and pressure gradients have on the deformation of neurons and the corresponding change in the flow pathway has not been addressed. While it is reasonable to assume a rigid flow channel network when the local pressure and pressure gradient are relatively low [54], it should be emphasised that (i) intermediate level of hydraulic pressure (e.g., 1500 Pa) can notably deform the axons and change the local permeability (Figs. 4 and 5); (ii) pressures higher than 1500 Pa widely exist in the brain as the normal ICP is 1000 ∼ 2000 Pa [20]. Under the conditions of hydrocephalus and intracranial hypertension, the local ICP can even reach 3332.5 Pa [55]. All the evidence underscores the importance of considering microstructural deformation when dealing with fluid/mass transport in the brain, which has been explored in this contribution.

How to model the development and dissipation of hydrocephalus is still an open question [56]; insufficient understanding of how the microstructure regulates the hydrops is one of the major obstacles. Looking at CED and other techniques aimed at circumventing issues linked to the presence of the blood brain barrier, which are regarded as promising treatments for multiple brain disorders [57, 58, 59], few clinical successes have been reported. This is also caused by the limited predictive capability of drug distribution in the brain tissue during such procedures due to the absence of pressure-permeability relationship that can be applied when modelling fluid transport in the brain, leading to most of the practical methods assuming a rigid flow channel network. We, therefore, believe that the microscale ISF-Axon interaction mechanism and the resulting macroscale pressure-dependent permeability tensor proposed in this study can, despite the inherent simplifications made to derive the permeability tensor using idealised geometries and descriptions of the complex biophysical interactions governing the problem, lay the groundwork for more advanced models to understand the pressure-related biophysical phenomena in the brain and also assist brain disorder treatments.

Poroelastic theory can also consider pressure-dependent permeability of porous media by treating the porous medium as a biphasic domain constituted by solid matrix and fluid phase [60]. In the control volume, an increase of the fluid occupation compresses the solid matrix and enlarges the porosity. By linking porosity to permeability by e.g. the well-known semi-empirical Kozeny-Carman formula, the relationship between flow status and permeability can be established [61]. Methods based on this theory have been widely applied in the brain-mechanics research [62]. According to our results (Figs. 4 and 5), these methods could be valid if the change of permeability is dominated by axons shrinkage, i.e., when the flow is parallel to axons. However, when displacements and contact of axons play an important role in affecting the fluid flow, i.e., when the flow is perpendicular to axons, the poroelastic theory is not capable of capturing all the decisive factors. To make it worse, the microstructural-dependence study (Section 3.4) has shown that the axon’s mechanical and geometrical properties also have a remarkable effect on WM’s equivalent permeability, which means that directly linking tissue porosity to tissue permeability without considering the axons’ deformation may lead to a significant loss of fidelity when using homogenisation techniques to describe fluid flow in the brain.

Results in Fig. 7 demonstrate the general compatibility of proposal ISF-Axon interaction theory in the ovine brain WM as different configurations of microstructure reached the same trend of pressure-permeability relationship. The emergence of high-resolution imaging techniques has made it possible to also obtain microstructural information of the human brain [63, 64], thus the protocol proposed in this study can be also used to gain a deeper understanding of ISF-Axon interaction and fluid/mass transport in the human brain. In addition, Figs. 6 and 7 also indicate the significance of microstructural properties (specifically the compressibility of axons, axon diameter, tissue porosity, and axon tortuosity) on the magnitude of permeability tensor and pressure-permeability relationship. As discussed above, the geometric information could be obtained by high-resolution imaging techniques, but the compressibility measurement of axons is still challenging, so no conclusion has been made on the exact value of Poisson’s ratio of axons. Although most of the mechanical studies acknowledged the near-incompressibility of axons, our results imply that it is worth to further explore the compressibility and other mechanical properties of neuronal cells and structures.

Many emerging technologies requires brain-tissue-like biomaterials to serve as brain organoids [65], scaffolds for neural tissue engineering [66], improve brain–machine interfaces [67] and surgical practise [21, 22]. While matching the mechanical property of the real brain to minimise immune response and implant rejection has been the major goal of many recent researchers, and accurate microstructural-informed constitutive models to capture brain mechanical behaviours have been developed by Holzapfel’s group and Kuhl’s group [68, 69] to support brain modelling and the design of these materials, the capability of mimicking brain’s hydraulic properties, such as poroviscoelastic responses coupled with fluid/mass transport property, has also been emphasised as an extremely important target by the advanced brain tissue engineering community [26]. In this study, apart from unrevealing the biophysical phenomena governing ISF-Axon interactions, the formulation of the newly proposed anisotropic pressure-dependent permeability tensor fills the gap by enabling us to capture the brain tissues’ complex hydraulic behaviours and tune the transport properties of brain-tissue-like biomaterials.

## 5. Conclusion

In the absence of advanced experimental techniques to characterise the effect of microscale ISF-Axon interaction on macroscale fluid transport in the brain, we have developed a representative geometric model of brain WM from Focused Ion-Beam Scanning Electron Microscopy (FIB-SEM) image data, and a comprehensive mathematical model to provide a better understanding of this complex multiscale biophysical phenomenon. An anisotropic pressure-dependent permeability tensor for brain WM was subsequently put forward and validated using the results from infusion experiments with ovine brain samples. Compared with poroelastic model, the newly developed permeability tensor can better capture the hydraulic behaviours and transport property of brain WM. As microstructural variance may lead to large difference of the permeability tensor, our study suggests it is critical to include microstructural information when characterising the transport properties of the brain. Our study, while providing tools to significantly improve the prediction of fluid/mass transport in brain and support the treatment of brain disorders, sets a blueprint for understanding the biophysical phenomena in other brain-like fibrous tissues, such as cartilage, muscle, and tendons to support various types of biomaterial design.

## Supplementary information

Movie S1: Dynamic response of the axons and corresponding change of the ISF flow status under parallel infusion. Pressure = 1600 Pa, Δ P = 12 Pa.

Movie S2: Dynamic response of the axons and corresponding change of the ISF flow status under perpendicular infusion. Pressure = 1600 Pa, Δ P = 3.5 Pa.

Movie S3: Axons’ deformation and ISF flow status after reaching equilibrium at different hydraulic pressure (10 ∼ 3000 Pa) under parallel infusion. The pressure drop did not change, which was 1 Pa. Movie S4: Axons’ deformation and ISF flow status after reaching equilibrium at different pressure drop (0.1 ∼ 12 Pa) under parallel infusion. The hydraulic pressure did not change, which was 1600 Pa.

Movie S5: Axons’ deformation and ISF flow status after reaching equilibrium at different hydraulic pressure (10 ∼ 3000 Pa) under perpendicular infusion. The pressure drop did not change, which was 1 Pa.

Movie S6: Axons’ deformation and ISF flow status after reaching equilibrium at different pressure drop (0.01 ∼ 5 Pa) under perpendicular infusion. The hydraulic pressure did not change, which was 1600 Pa.

## Acknowledgements

This project has received funding from the European Unions Horizon 2020 research and innovation programme under Grant Agreement No. 688279. Daniele Dini would like to acknowledge the support received from the EP-SRC under the Established Career Fellowship Grant No. EP/N025954/1. Wenbo Zhan would like to acknowledge the support received from the Children with Cancer UK under the project Children’s Brain Tumour Drug Delivery Consortium Grant No. 16-224.

## CRediT authorship contribution statement

**Tian Yuan:** Conceptualisation, Methodology, Software, Writing - Original draft preparation. **Wenbo Zhan:** Methodology, Discussion, Writing, Reviewing. **Daniele Dini:** Conceptualisation, Supervision, Reviewing.

## References

[1] M. Ito, Control of mental activities by internal models in the cerebellum, Nat. Rev. Neurosci. 9 (2008) 304–313. doi:10.1038/nrn2332.

[2] A. D. Craig, Interoception: the sense of the physiological condition of the body, Curr. Opin. Neurobiol. 13 (4) (2003) 500–505. doi: 10.1016/S0959-4388(03)00090-4.

[3] S. B. Hladky, M. A. Barrand, Mechanisms of fluid movement into, through and out of the brain: evaluation of the evidence, Fluids Barriers CNS 11 (1) (2014) 1–32. doi:10.1186/2045-8118-11-26.

[4] M. D. Sweeney, Z. Zhao, A. Montagne, A. R. Nelson, B. V. Zlokovic, Blood-brain barrier: From physiology to disease and back, Physiol. Rev. 99 (1) (2019) 21–78. doi:10.1152/physrev.00050.2017.

[5] L. Xie, H. Kang, Q. Xu, M. J. Chen, Y. Liao, M. Thiyagarajan, J. O’Donnell, D. J. Christensen, C. Nicholson, J. J. Iliff, T. Takano, R. Deane, M. Nedergaard, Sleep drives metabolite clearance from the adult brain, Science 342 (6156) (2013) 373–377. doi:10.1126/science.1241224.

[6] D. D. Shen, A. A. Artru, K. K. Adkison, Principles and applicability of csf sampling for the assessment of cns drug delivery and pharmacodynamics, Adv. Drug Delivery Rev. 56 (12) (2004) 1825–1857. doi:10.1016/j.addr.2004.07.011.

[7] K. Sawamoto, H. Wichterle, O. Gonzalez-Perez, J. A. Cholfin, M. Yamada, N. Spassky, N. S. Murcia, J. M. Garcia-Verdugo, O. Marin, J. L. R. Rubenstein, M. Tessier-Lavigne, H. Okano, A. Alvarez-Buylla, New neurons follow the flow of cerebrospinal fluid in the adult brain, Science 311 (5761) (2006) 629–632. doi:10.1126/science.1119133.

[8] L. D. Lewis, The interconnected causes and consequences of sleep in the brain, Science 374 (6567) (2021) 564–568. doi:10.1126/science.abi8375.

[9] N. E. Fultz, G. Bonmassar, K. Setsompop, R. A. Stickgold, B. R. Rosen, J. R. Polimeni, L. D. Lewis, Coupled electrophysiological, hemodynamic, and cerebrospinal fluid oscillations in human sleep, Science 366 (6465) (2019) 628–631. doi:10.1126/science.aax5440.

[10] P. Brundin, R. Melki, R. Kopito, Prion-like transmission of protein aggregates in neurodegenerative diseases, Nat. Rev. Mol. Cell Biol. 11 (4) (2010) 301. doi:10.1038/nrm2873.

[11] J. Weickenmeier, E. Kuhl, A. Goriely, Multiphysics of prionlike diseases: Progression and atrophy, Phys. Rev. Lett. 121 (15) (2018) 158101. doi:10.1103/PhysRevLett.121.158101.

[12] H. Bobo R, W. Laske D, A. Akbasak, F. Morrison P, L. Dedrick R, H. Oldfield E, Convection-enhanced delivery of macromolecules in the brain., Proc. Natl. Acad. Sci. U.S.A. 91 (6) (1994) 2076–2080. doi:10.1073/pnas.91.6.2076.

[13] A. Jamal, T. Yuan, S. Galvan, A. Castellano, M. Riva, R. Secoli, A. Falini, L. Bello, F. Rodriguez y. Baena, D. Dini, Insights into Infusion-Based Targeted Drug Delivery in the Brain: Perspectives, Challenges and Opportunities, Int. J. Mol. Sci. 23 (6) (2022) 3139. doi:10.3390/ijms23063139.

[14] M. K. Rasmussen, H. Mestre, M. Nedergaard, Fluid transport in thebrain, Physiol. Rev. 102 (2) (2022) 1025–1151. doi:10.1152/physrev.00031.2020.

[15] H. K. Erik, K. Benjamin, D. Anna, J. Sejnowski Terrence, M. Dale Anders, W. Omholt Stig, O. O. Petter, N. E. Arnulf, M. Kent-André, H. Pettersen Klas, Interstitial solute transport in 3d reconstructed neuropil occurs by diffusion rather than bulk flow, Proc. Natl. Acad. Sci. U.S.A. 114 (37) (2017) 9894–9899. doi:10.1073/pnas.1706942114.

[16] A.-R. A. Khaled, K. Vafai, The role of porous media in modelling flow and heat transfer in biological tissues, Int. J. Heat Mass Transfer 46 (26) (2003) 4989–5003. doi:10.1016/S0017-9310(03)00301-6.

[17] T. L. Chenevert, J. A. Brunberg, J. G. Pipe, Anisotropic diffusion in human white matter: demonstration with mr techniques in vivo., Radiology 177 (2) (Nov 1990). doi:10.1148/radiology.177.2.2217776.

[18] M. Vidotto, A. Bernardini, M. Trovatelli, E. De Momi, D. Daniele, On the microstructural origin of brain white matter hydraulic permeability, Proc. Natl. Acad. Sci. U.S.A. 118 (36) (2021) e2105328118. doi:10.1073/pnas.2105328118.

[19] L. Yun-Bi, F. Kristian, S. Gerald, S. Christian, K. Frank, W. Hartwig, G. Jochen, J. Paul, W. Er-Qing, K. Josef, R. Andreas, Viscoelastic properties of individual glial cells and neurons in the cns, Proc. Natl. Acad. Sci. U.S.A. 103 (47) (2006) 17759–17764. doi:10.1073/pnas.0606150103.

[20] D. Renier, C. Sainte-Rose, D. Marchac, Intracranial pressure in cran-iostenoses, in: Craniofacial Surgery, Springer, Berlin, Germany, 1987, pp. 110–113. doi:10.1007/978-3-642-82875-1_24.

[21] Z. Tan, D. Dini, F. Rodriguez y. Baena, A. E. Forte, Composite hydrogel: A high fidelity soft tissue mimic for surgery, Mater. Des. 160 (2018) 886–894. doi:10.1016/j.matdes.2018.10.018.

[22] A. E. Forte, S. Galvan, F. Manieri, F. Rodriguez y. Baena, D. Dini, A composite hydrogel for brain tissue phantoms, Mater. Des. 112 (2016) 227–238. doi:10.1016/j.matdes.2016.09.063.

[23] A. E. Forte, S. M. Gentleman, D. Dini, On the characterization of the heterogeneous mechanical response of human brain tissue, Biomech. Model. Mechanobiol. 16 (3) (2017) 907–920. doi:10.1007/s10237-016-0860-8.

[24] A. Dine, E. Bentley, L. A. PoulmarcK, D. Dini, A. E. Forte, Z. Tan, A dual nozzle 3D printing system for super soft composite hydrogels, HardwareX 9 (2021) e00176. doi:10.1016/j.ohx.2021.e00176.

[25] M. Terzano, A. Spagnoli, D. Dini, A. E. Forte, Fluid–solid interaction in the rate-dependent failure of brain tissue and biomimicking gels, J. Mech. Behav. Biomed. Mater. 119 (2021) 104530. doi:10.1016/j.jmbbm.2021.104530.

[26] E. Axpe, G. Orive, K. Franze, E. A. Appel, Towards brain-tissue-like biomaterials, Nat. Commun. 11 (3423) (2020) 1–4. doi:10.1038/s41467-020-17245-x.

[27] A. Jamal, M. T. Mongelli, M. Vidotto, M. Madekurozwa, A. Bernardini, D. R. Overby, E. De Momi, F. R. y. Baena, J. M. Sherwood, D. Dini, Infusion mechanisms in brain white matter and their dependence on microstructure: An experimental study of hydraulic permeability, IEEE Trans. Biomed. Eng. 68 (4) (2020) 1229–1237. doi:10.1109/TBME.2020.3024117.

[28] B. Tully, Y. Ventikos, Cerebral water transport using multiple-network poroelastic theory: application to normal pressure hydrocephalus, J. Fluid Mech. 667 (2011) 188–215. doi:10.1017/S0022112010004428.

[29] L. Guo, J. C. Vardakis, D. Chou, Y. Ventikos, A multiple-network poroelastic model for biological systems and application to subject-specific modelling of cerebral fluid transport, Int. J. Eng. Sci. 147 (2020) 103204. doi:10.1016/j.ijengsci.2019.103204.

[30] A. Goriely, M. G. Geers, G. A. Holzapfel, J. Jayamohan, A. Jérusalem, S. Sivaloganathan, W. Squier, J. A. van Dommelen, S. Waters, E. Kuhl, Mechanics of the brain: perspectives, challenges, and opportunities, Biomechanics and modeling in mechanobiology 14 (5) (2015) 931–965.

[31] S. Budday, P. Bayly, G. A. Holzapfel, Advances in brain mechanics, Frontiers in Mechanical Engineering (2021) 106.

[32] W. Yumei, W. Christina, X. C. Shan, J. Hayworth Kenneth, J. Weinberg Richard, F. Hess Harald, D. C. Pietro, Contacts between the endoplasmic reticulum and other membranes in neurons, Proc. Natl. Acad. Sci. U.S.A. 114 (24) (2017) E4859–E4867. doi:10.1073/pnas.1701078114.

[33] A. I. Silva, J. E. Haddon, Y. A. Syed, S. Trent, T.-C. E. Lin, Y. Patel, J. Carter, N. Haan, R. C. Honey, T. Humby, Y. Assaf, M. J. Owen, D. E. J. Linden, J. Hall, L. S. Wilkinson, Cyfip1 haploinsufficient rats show white matter changes, myelin thinning, abnormal oligodendrocytes and behavioural inflexibility, Nat. Commun. 10 (1) (2019) 3455. arXiv:31371763, doi:10.1038/s41467-019-11119-7.

[34] A. Bernardini, M. Trovatelli, M. M. Kłosowski, M. Pederzani, D. D. Zani, S. Brizzola, A. Porter, F. Rodriguez y. Baena, D. Dini, Reconstruction of ovine axonal cytoarchitecture enables more accurate models of brain biomechanics, Commun. Biol. 5 (1101) (2022) 1–14. doi:10.1038/s42003-022-04052-x.

[35] E. Syková, C. Nicholson, Diffusion in brain extracellular space, Physiol. Rev. 88 (4) (Oct 2008).

[36] S. A. Yousefsani, F. Farahmand, A. Shamloo, A three-dimensional micromechanical model of brain white matter with histology-informed probabilistic distribution of axonal fibers, J. Mech. Behav. Biomed. Mater. 88 (2018) 288–295. doi:10.1016/j.jmbbm.2018.08.042.

[37] J. H. Kim, G. W. Astary, S. Kantorovich, T. H. Mareci, P. R. Carney, M. Sarntinoranont, Voxelized computational model for convection-enhanced delivery in the rat ventral hippocampus: comparison with in vivo mr experimental studies, Ann. Biomed. Eng. 40 (9) (2012) 2043–2058. arXiv:22532321, doi:10.1007/s10439-012-0566-8.

[38] W. Dai, G. W. Astary, A. K. Kasinadhuni, P. R. Carney, T. H. Mareci, M. Sarntinoranont, Voxelized model of brain infusion that accounts for small feature fissures: Comparison with magnetic resonance tracer studies, J. Biomech. Eng. 138 (5) (2016) 051007. arXiv:26833078, doi:10.1115/1.4032626.

[39] T. Yuan, L. Gao, W. Zhan, D. Dini, Effect of particle size and surface charge on nanoparticles diffusion in the brain white matter, Pharm. Res. (2022) 1–15doi:10.1007/s11095-022-03222-0.

[40] T. Yuan, W. Zhan, A. Jamal, D. Dini, On the microstructurally driven heterogeneous response of brain white matter to drug infusion pressure, Biomechanics and Modeling in Mechanobiology 21 (4) (2022) 1299–1316.

[41] L. M. Wang, E. Kuhl, Viscoelasticity of the axon limits stretch-mediated growth, Comput. Mech. 65 (3) (2020) 587–595. doi:10.1007/s00466-019-01784-2.

[42] R. Bernal, P. A. Pullarkat, F. Melo, Mechanical properties of axons, Phys. Rev. Lett. 99 (1) (2007) 018301. doi:10.1103/PhysRevLett.99.018301.

[43] L. Guo, Y. Wang, High-rate tensile behavior of silicone rubber at various temperatures, Rubber Chem. Technol. 93 (1) (2020) 183–194. doi:10.5254/rct.19.81562.

[44] D. F. Meaney, Relationship between structural modeling and hyperelastic material behavior: application to cns white matter, Biomech. Model. Mechanobiol. 1 (4) (2003) 279–293. doi:10.1007/s10237-002-0020-1.

[45] COMSOL-Multiphysics, The comsol multiphysics reference manual, v5.6 (2020). URL http://www.comsol.com/products/multiphysics/

[46] Y. Zhang, K. Abiraman, H. Li, D. M. Pierce, A. V. Tzingounis, G. Lykotrafitis, Modeling of the axon membrane skeleton structure and implications for its mechanical properties, PLoS Comput. Biol. 13 (2) (2017) e1005407. doi:10.1371/journal.pcbi.1005407.

[47] J. A. Galbraith, L. E. Thibault, D. R. Matteson, Mechanical and electrical responses of the squid giant axon to simple elongation, J. Biomech. Eng. 115 (1) (1993) 13–22. doi:10.1115/1.2895464.

[48] T. L. Haut Donahue, M. L. Hull, M. M. Rashid, C. R. Jacobs, How the stiffness of meniscal attachments and meniscal material properties affect tibio-femoral contact pressure computed using a validated finite element model of the human knee joint, J. Biomech. 36 (1) (2003) 19–34. doi:10.1016/S0021-9290(02)00305-6.

[49] S. Behforootan, P. E. Chatzistergos, N. Chockalingam, R. Naemi, A clinically applicable non-invasive method to quantitatively assess the visco-hyperelastic properties of human heel pad, implications for assessing the risk of mechanical trauma, J. Mech. Behav. Biomed. Mater. 68 (2017) 287–295. doi:10.1016/j.jmbbm.2017.02.011.

[50] A. Samadi-Dooki, G. Z. Voyiadjis, R. W. Stout, A combined experimental, modeling, and computational approach to interpret the viscoelastic response of the white matter brain tissue during indentation, J. Mech. Behav. Biomed. Mater. 77 (2018) 24–33. doi:10.1016/j.jmbbm.2017.08.037.

[51] S. A. Yousefsani, A. Shamloo, F. Farahmand, Micromechanics of brain white matter tissue: A fiber-reinforced hyperelastic model using embedded element technique, J. Mech. Behav. Biomed. Mater. 80 (2018) 194–202. doi:10.1016/j.jmbbm.2018.02.002.

[52] M. Vidotto, D. Botnariuc, E. De Momi, D. Dini, A computational fluid dynamics approach to determine white matter permeability, Biomech. Model. Mechanobiol. 18 (4) (2019) 1111. doi:10.1007/s10237-019-01131-7.

[53] D. S. Clague, B. D. Kandhai, R. Zhang, P. M. A. Sloot, Hydraulic per-meability of (un)bounded fibrous media using the lattice boltzmann method, Phys. Rev. E 61 (1) (2000) 616–625. doi:10.1103/PhysRevE.61.616.

[54] A. A. Linninger, M. R. Somayaji, T. Erickson, X. Guo, R. D. Penn, Computational methods for predicting drug transport in anisotropic and heterogeneous brain tissue, J. Biomech. 41 (10) (2008) 2176–2187. doi:10.1016/j.jbiomech.2008.04.025.

[55] M. Czosnyka, J. D. Pickard, Monitoring and interpretation of intracranial pressure, J. Neurol. Neurosurg. Psychiatry 75 (6) (2004) 813–821. doi:10.1136/jnnp.2003.033126.

[56] S. Budday, T. C. Ovaert, G. A. Holzapfel, P. Steinmann, E. Kuhl, Fifty shades of brain: A review on the mechanical testing and modeling of brain tissue, Arch. Comput. Methods Eng. 27 (4) (2020) 1187–1230. doi:10.1007/s11831-019-09352-w.

[57] W. Debinski, S. B. Tatter, Convection-enhanced delivery for the treatment of brain tumors, Expert Rev. Neurother. 9 (10) (2009) 1519–1527. doi:10.1586/ern.09.99.

[58] M. A. Rogawski, Convection-enhanced delivery in the treatment of epilepsy, Neurotherapeutics 6 (2) (2009) 344–351. doi:10.1016/j.nurt.2009.01.017.

[59] M. S. Fiandaca, J. R. Forsayeth, P. J. Dickinson, K. S. Bankiewicz, Image-guided convection-enhanced delivery platform in the treatment of neurological diseases, Neurotherapeutics 5 (1) (2008) 123–127. doi:10.1016/j.nurt.2007.10.064.

[60] A. H.-D. Cheng, Poroelasticity, Springer, Cham, Switzerland, 2016. doi:10.1007/978-3-319-25202-5.

[61] C. W. MacMinn, E. R. Dufresne, J. S. Wettlaufer, Large deformations of a soft porous material, Phys. Rev. Appl. 5 (4) (2016) 044020. doi:10.1103/PhysRevApplied.5.044020.

[62] A. Goriely, M. G. D. Geers, G. A. Holzapfel, J. Jayamohan, A. Jérusalem, S. Sivaloganathan, W. Squier, J. A. W. van Dommelen, S. Waters, E. Kuhl, Mechanics of the brain: perspectives, challenges, and opportunities, Biomech. Model. Mechanobiol. 14 (5) (2015) 931–965. doi:10.1007/s10237-015-0662-4.

[63] L. J. Edwards, E. Kirilina, S. Mohammadi, N. Weiskopf, Microstructural imaging of human neocortex in vivo, Neuroimage 182 (2018) 184–206. doi:10.1016/j.neuroimage.2018.02.055.

[64] E. Nishat, S. Stojanovski, S. E. Scratch, S. H. Ameis, A. L. Wheeler, Premature white matter microstructure in female children with a history of concussion, medRxiv (2021) 2021.12.02.21267220arXiv: 2021.12.02.21267220.

[65] M. A. Lancaster, M. Renner, C.-A. Martin, D. Wenzel, L. S. Bicknell, M. E. Hurles, T. Homfray, J. M. Penninger, A. P. Jackson, J. A. Knoblich, Cerebral organoids model human brain development and microcephaly, Nature 501 (2013) 373–379. doi:10.1038/nature12517.

[66] F. Li, M. Ducker, B. Sun, F. G. Szele, J. T. Czernuszka, Interpenetrating polymer networks of collagen, hyaluronic acid, and chondroitin sulfate as scaffolds for brain tissue engineering, Acta Biomater. 112 (2020) 122–135. doi:10.1016/j.actbio.2020.05.042.

[67] T. D. Kozai, K. Catt, X. Li, Z. V. Gugel, V. T. Olafsson, A. L. Vazquez, X. T. Cui, Mechanical failure modes of chronically implanted planar silicon-based neural probes for laminar recording, Biomaterials 37 (2015) 25–39.

[68] S. Budday, M. Sarem, L. Starck, G. Sommer, J. Pfefferle, N. Phunchago, E. Kuhl, F. Paulsen, P. Steinmann, V. P. Shastri, G. A. Holzapfel, Towards microstructure-informed material models for human brain tissue, Acta Biomater. 104 (2020) 53–65. doi:10.1016/j.actbio.2019.12.030.

[69] M. Hoppstädter, D. Püllmann, R. Seydewitz, E. Kuhl, M. Böl, Correlating the microstructural architecture and macrostructural behaviour of the brain, Acta Biomater. 151 (2022) 379–395. doi:10.1016/j.actbio.2022.08.034.

